# DNA methylation and gene expression dynamics during cotton ovule and fiber development

**DOI:** 10.1101/010702

**Authors:** Qingxin Song, Xueying Guan, Z. Jeffrey Chen

## Abstract

Cotton is the largest source of renewable textile fiber and a successful model of transgenic applications in crop production. However, improving cotton production using fiber-related transgenes is somewhat difficult. This is probably related to unique epigenetic and gene expression changes during fiber development. Here we show that inhibiting DNA methylation impairs fiber development. Genome-wide methylcytosine-, mRNA-, and small RNA-sequencing analyses reveal minor changes in CG and CHG methylation and distinct changes in CHH methylation among different tissues. In ovules CHH hypermethyaltion is associated with small RNA-directed DNA methylation (RdDM) and expression changes of nearby genes in euchromatin. Remarkably, ovule-derived fiber cells not only maintain euchromatic CHH methylation, but also generate additional heterochromatic CHH hypermethylation independent of RdDM, which represses transposable elements (TEs) and nearby genes including fiber-related genes. Furthermore, DNA methylation contributes to the expression bias of homoeologous genes in ovules and fibers. This spatiotemporal DNA methylation in promoters could act as a double-lock feedback mechanism to regulate TE and gene expression, which could be translated into genomic and biotechnological improvement of agronomic traits.

---

The most widely-cultivated cotton (*Gossypium hirsutum* L., AADD) is an allotetraploid species, which originated 1-2 million years ago from interspecific hybridization between A-genome species, resembling *Gossypium herbaceum* or *Gossypium arboretum*, and D-genome species, resembling *Gossypium raimondii* (Wendel 1989). The intergenomic interaction in allotetraploid cottons induces longer fiber and higher yield, coincident with expression bias of fiber-related homoeologous genes (Flagel et al. 2008; Pang et al. 2009), which provides the basis of selection and domestication for agronomic traits in cotton and many other polyploid crops (Wendel 1989; Chen et al. 2007). Each cotton fiber is a singular cell derived from an epidermal cell of the ovule, undergoing rapid cell elongation and cellulose biosynthesis, and ˜100,000 fiber cells develop semi-synchronically in each ovule (seed) and can reach six centimeters in length (Kim and Triplett 2001; Guan et al. 2014). In early stages of fiber development, rapid cell growth is associated with a dramatic increase of DNA content by endoreduplication (Van’t Hof 1999; Wu et al. 2006) and dynamic changes in gene expression and small RNAs (Pang et al. 2009; Guan et al. 2014; Yoo and Wendel 2014). Interestingly, DNA methylation changes are related to seasonal variation of fiber development in cotton (Jin et al. 2013) and is also shown to change among different tissues including fibers based on the methylation-sensitive high-performance liquid chromatography (HPLC) assay (Osabe et al. 2014). Moreover, over-expressing fiber-related transgenes often leads to the unexpected outcome of fiber phenotypes (Walford et al. 2011). These data indicate a potential role for DNA methylation in gene expression and phenotypic traits such as cotton fiber, which could be selected by breeding.

DNA methylation is heritable epigenetic information in most eukaryotes and mediates many epigenetic phenomena, including imprinting and transposon silencing (Bird 1992; Richards 1997; Law and Jacobsen 2010; Haag and Pikaard 2011; Schultz et al. 2012). In plants, DNA is methylated in CG, CHG and CHH (H=A, T, or C) sites through distinct pathways. In *Arabidopsis*, CG methylation is maintained by METHYLTRANSFERASE1 (MET1) (Kankel et al. 2003), a homolog of mammalian DNMT1 (Li et al. 1992). Plant-specific CHROMOMETHYLASE3 (CMT3) is primarily responsible for CHG methylation, which is coupled with H3K9 dimethylation (Lindroth et al. 2001; Cao and Jacobsen 2002). CHH methylation is established de novo by DOMAINS REARRANGED METHYLTRANSFERASE1 and 2 (DRM1 and DRM2) (Cao et al. 2003) through the RNA-directed DNA methylation (RdDM) pathway (Wassenegger et al. 1994), involving 24-nt small interfering RNAs (siRNAs) (Law and Jacobsen 2010; Haag and Pikaard 2011). Recent studies found that independent of RdDM, CHH methylation could also be established by CMT2 (Zemach et al. 2013; Stroud et al. 2014), through histone H1 and DECREASE-IN-DNA-METHYLATION1 (DDM1) activities (Jeddeloh et al. 1999). The methylome data indicate that CMT2 and RdDM pathways preferentially function in heterochromatic and euchromatic regions, respectively (Zemach et al. 2013; Stroud et al. 2014).

Although genome-wide DNA methylation has been examined in Arabidopsis (Zhang et al. 2006), soybean (Song et al. 2013), maize (Gent et al. 2013), and other plants and animals (Zemach et al. 2010), the biological roles of RdDM and CMT2-depednent methylation pathways remain elusive. Moreover, DNA methylation patterns and changes during cotton fiber development are unknown. In this study, we tested the effects of DNA methylation on cotton ovule and fiber development. Using methylcytosine-sequencing (MethylC-seq), RNA-seq, and small RNA-seq analyses, we examined CG, CHG, and CHH methylation patterns in fibers, ovules, and leaves and analyzed differentially methylated regions (DMRs) between the ovule and leaf (OL) and between the fiber and ovule (FO). The methylation patterns in the gene body and 5’ and 3’ flanking sequences were comparatively analyzed with TE densities and expression levels of nearby genes as well as small RNA loci. The results provide unique roles of CG, CHG, and CHH methylation in cotton ovule and fiber development and biased expression of homoeologous genes.

## RESULTS

### DNA methylation dynamics in cotton leaves, ovules, and fibers

DNA methylation affects growth and development in plants and animals (Bird 1992; Richards 1997). Methylation levels change among different cotton tissues (Osabe et al. 2014). To investigate the biological role of DNA methylation in fiber development, we harvested ovules at -3-0 days post anthesis (DPA) from the allotetraploid cotton Gossypium hirsutum L. acc. TM-1 and cultured them in vitro with or without treatment of the DNA methyltransferase inhibitor, 5-aza-deoxycytidine (5-aza-dC) (Haaf et al. 1993). Fiber length and ovule size were suppressed in the initiation and elongation stages by 5-aza-dC treatment (Fig. 1A-C), providing pharmacological evidence for the requirement of DNA methylation during fiber and ovule development.

**Figure 1.**
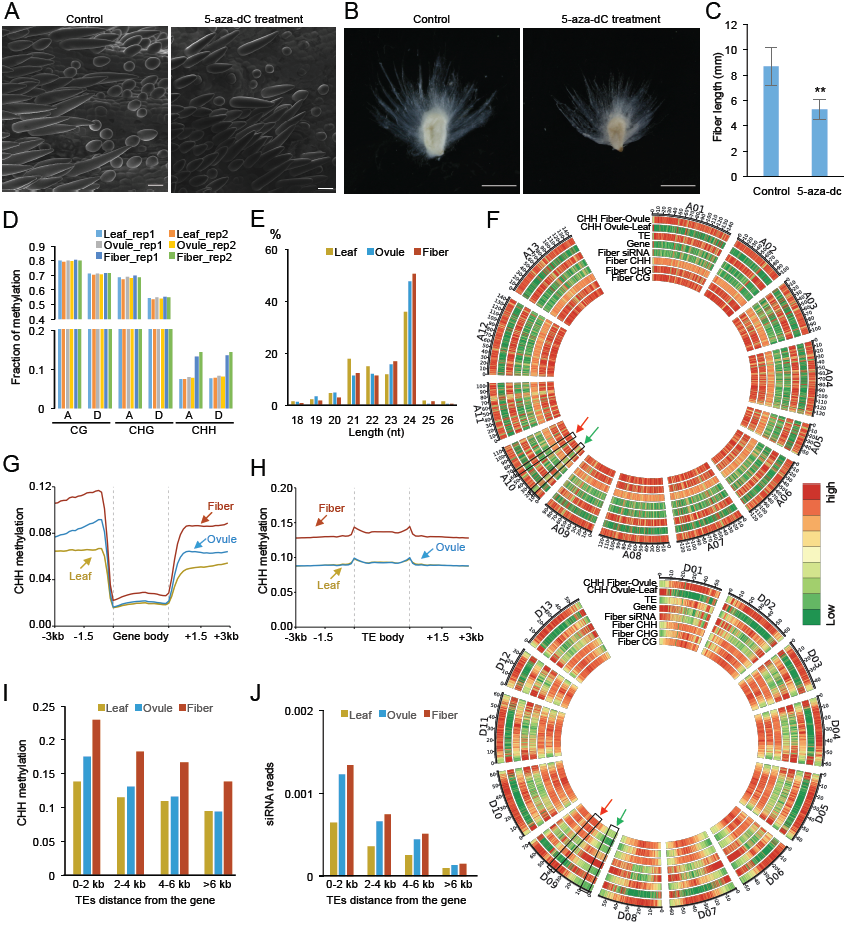
Effects of DNA methylation on ovule and fiber development and genome-wide distribution of DNA methylation in leaves, ovules, and fibers. (*A*) Scanning electron micrograph (SEM) showing shorter and fewer fibers in ovules treated by 5-aza-dC. Ovules at -3--4 DPA were cultured in vitro for 4 days with and without 5-aza-dC (10mg/L). (*B*) Shorter fibers in ovules after the aza-dC treatment for 14 days. (*C*) Quantitative analysis of fiber length in (*B*) (student’s test, p<0.01). (*D*) Percentage of methylated cytosine (mC) in leaves, ovules, and fibers. A and D indicate A- and D-subgenomes in the allotetraploid cotton, respectively. (*E*) Size distribution of small RNAs in different tissues. (*F*) Circle plots of fiber CG, CHG, CHH methylation, fiber small RNA density, gene density, TE density, and ovule-leaf and fiber-ovule CHH methylation among 12 A-homoeologous chromosomes (upper) and 13 D-homoeologous chromosomes (lower). The color bar indicates the same scale that was used from low (green) to high (red) densities. (*G*) Distribution of CHH methylation in genes. (*H*) Distribution of CHH methylation in TEs. (*I*) CHH methylation levels in TEs relative to the distance from the nearest gene. (*J*) siRNA levels (per bp per million reads) in TEs relative to the distance from the nearest gene.

This result led to the investigation of genome-wide DNA methylation changes during ovule and fiber development. We performed whole-genome bisulfite sequencing in leaves, ovules at 0 DPA, and fibers at 14 DPA with two biological replicates and ovules at 14 DPA with one biological replicate. The genomes of G. arboreum (Li et al. 2014) and *G. raimondii* (Paterson et al. 2012) were modified using single nucleotide polymorphism (SNP) generating from genome sequencing reads of *G. hirsutum*, and then used as references for mapping sequence reads and data analysis. The bisulfite-conversion rates were over 99% (Supplemental Table S1). Approximately 80% of cytosines were covered by at least one uniquely mapped read, and the mean coverage of cytosines was 7.5-fold or higher for all tissues. Methylation levels between biological replicates were highly correlated (Pearson r>0.8), indicating reproducibility of the data (Fig. 1D; Supplemental Fig. S1A). Overall, CG and CHG methylation levels were similar among fibers, ovules and leaves but were lower in the D-subgenome than in the A-subgenome for all tissues (Fig. 1D). However, CHH methylation levels were much higher in fibers (˜14%) than in ovules (˜8.1%) and leaves (˜7.6%) and were slightly higher in the D-subgenome than in the A-genome for all tissues (Fig. 1D). This high methylation level in fibers was not related to the developmental stage because the methylation level in ovules was slightly lower at 14 DPA than at 0 DPA (Supplemental Fig. S1B). To study functions of small RNAs in DNA methylation changes in cotton tissues, we generated small RNA-seq data using leaves, ovules at 0 DPA and fibers at 14 DPA. The high CHH methylation level in fibers correlated with the small RNA-seq data, in which 24-nt siRNAs were higher in fibers and ovules than in leaves (Fig. 1E) and consistent with the published small RNA-seq data (Pang et al. 2009).

On each chromosome, DNA methylation was more abundant in transposable element (TE)-rich than gene-rich regions in all tissues (Fig. 1F; Supplemental Fig. S1C). However, 24-nt siRNAs, which induce CHH methylation through RdDM (Haag and Pikaard 2011; Law et al. 2013), were highly enriched in gene-rich regions. These 24-nt siRNAs were derived from TEs but not from coding sequences in gene-rich regions (Supplemental Fig. S1D). Because each fiber cell is expanded from an epidermal cell of the ovule, further comparison was made between the ovule and leaf (OL) and/or between the fiber and ovule (FO), which represented the transitional stages between ovule and fiber development. OL CHH hypermethylation correlated with gene-rich regions, showing the same trend as the siRNAs (Fig. 1F). On the contrary, FO CHH hypermethylation was enriched in TE- and repeat-rich regions, characteristic of heterochromatin. The data indicate that OL CHH hypermethylation preferentially occurs in euchromatic regions, whereas FO CHH hypermethylation predominates in heterochromatic regions (Fig. 1F; Supplemental Fig. S1E).

While CG and CHG methylation levels in genes and TEs were similar among different tissues (Supplemental Fig. S2), CHH methylation differences in genes were primarily distributed in 5’ upstream and 3’ downstream sequences, especially between the ovule and leaf (Fig. 1G). Among all TEs, mean CHH methylation levels were higher in the fiber but indistinguishable between the ovule and leaf (Fig. 1H). The higher CHH methylation levels in fibers and ovules were correlated with the closer distances of TEs to the gene (Fig. 1I). When TEs were more than 4-6-kb away from the gene, CHH methylation levels became similar between the ovule and leaf. Consistently, the CHH methylation levels mirrored the siRNA distribution patterns, which decreased as they were further away from the gene, indicating the role of the RdDM pathway in CHH methylation of TEs near the gene (Fig. 1J).

**Differentially methylated regions**

As CG and CHG methylation levels were similar among different tissues examined, further analysis was focused on CHH methylation changes among these tissues. We predict that differentially methylated regions (DMRs) between tissues could play a role in biological function. To test this, we identified 28,108 CHH-hypermethylated DMRs between the ovule and leaf (OL CHH-hyper DMRs) and 94,449 CHH-hypermethylated DMRs between the fiber and ovule (FO CHH-hyper DMRs) (Supplemental Table S2). A subset of these DMRs was validated by bisulfite-sequencing individually cloned genomic fragments (Supplemental Fig. S3), confirming the results of these DMRs from genome-wide analysis. Compared to genome-wide distributions, both OL and FO CHH-hyper DMRs were under-represented in the gene-body (especially exon) (Fig. 2A). However, OL CHH-hyper DMRs predominated in intergenic regions (73%) relative to the genome-wide average (30%), whereas FO CHH-hyper DMRs were enriched in TEs (69% vs. 60%). Among TEs, OL CHH-hyper DMRs were more abundant in TEs <1-kb (68%) than the genome-wide average (30%), while FO CHH-hyper DMR distribution was similar to the genome-wide distribution (Fig. 2B). Methylation levels of most OL CHH-hyper DMRs were higher in fibers than in ovules, indicating that OL CHH-hyper DMRs continued to be hypermethylated in elongating fibers after they elongated from ovule epidermal cells (Fig. 2C). Interestingly, among the OL CHH-hyper DMRs, siRNA expression levels were significantly higher in ovules than in leaves (P < 1e^-100^, Wilcoxon rank-sum test), and this difference was not obvious between the fiber and ovule in FO CHH-hyper DMRs (Fig. 2D). The data suggest that siRNAs induce CHH hypermethylation in the ovule, whereas the CHH methylation increase in the fiber was independent of the RdDM pathway. In addition to CHH methylation, CG and CHG methylation levels also increased in both FO CHH-hyper DMRs and OL CHH-hyper DMRs, and CHG methylation increased more than CG methylation (Fig. 2E).

**Figure 2.**
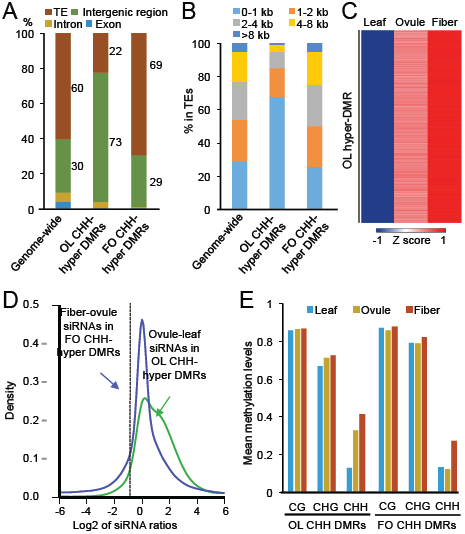
Genomic distribution of differentially methylated regions (DMRs). (*A*) Percentage of DMRs in TE (brown), intron (dark yellow), exon (blue), and intergenic region (green). (*B*) Percentage of DMRs in TEs with different sizes. (*C*) Heatmap of CHH methylation levels in OL CHH-hyper DMRs in leaves, ovules and fibers. (*D*) Kernel density plot of 24-nt siRNA fold change (ovule vs. leaf or fiber vs. ovule) in OL CHH-hyper DMRs and FO CHH-hyper DMRs. (*E*) Percentage of CG, CHG and CHH methylation in OL CHH-hyper DMRs and FO CHH-hyper DMRs.

### Effects of OL CHH-hyper DMR on gene expression

To test the role of methylation changes in gene expression, we compared RNA-seq data with DMRs between the ovule and leaf and/or between the fiber and ovule. Down-regulated genes in the ovule relative to the leaf were significantly enriched in the genes overlapping with OL CHH-hyper DMRs in the gene body (Fig. 3A; Supplemental Table S3), which is consistent with the notion that CHH methylation in the gene body correlates negatively with gene expression (Schmitz et al. 2013). However, up-regulated genes in ovules were significantly enriched in the genes overlapped with OL CHH-hyper DMRs in the flanking sequences (1 kb), especially upstream sequences (Fig. 3A; Supplemental Table S3), indicating a positive correlation of CHH methylation in flanking sequence with gene expression. For the genes that were preferentially expressed in the ovule (ovule-preferred genes), the CHH methylation level in the flanking sequences was higher in the ovule than in the leaf, whereas CHH methylation in flanking sequences of the leaf-preferred genes was similar between the ovule and leaf (Fig. 3B). To further test the relationship between CHH methylation and gene expression, we divided genes into quartiles based on their expression levels in each tissue (from low to high) and compared them with DNA methylation differences. CHH and CHG methylation in the gene body were negatively associated with gene expression, and moderately transcribed genes showed the highest CG methylation level in the gene body (Fig. 3C; Supplemental Fig. S4), which are consistent with previous findings (Zilberman et al. 2007; Schmitz et al. 2013). Moreover, CHH methylation in the 5’ upstream coincided with siRNA abundance, which was positively associated with gene expression (Fig. 3C,D). The siRNA levels in the promoter regions of ovule-preferred genes were significantly higher in ovules than in leaves (P < 1e^-4^, Wilcoxon rank-sum test) (Fig. 3E). These results indicate a unique role for the RdDM pathway in OL CHH hypermethylation and up-regulation of genes in ovules. This positive correlation between CHH methylation and gene expression was unexpected but also reported in other plants including maize (Gent et al. 2013) and soybean (Song et al. 2013).

**Figure 3.**
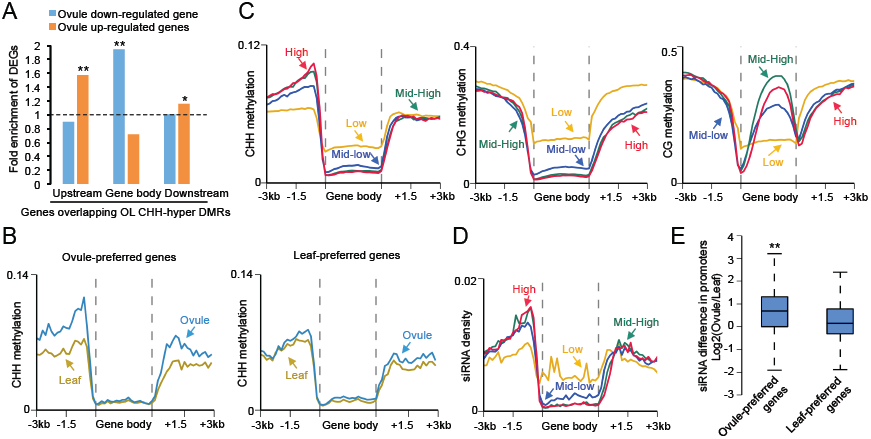
Relationship of ovule-leaf (OL) CHH-hyper DMRs and gene expression changes between the ovule and leaf. (*A*) Fold-enrichment of differentially expressed genes (DEGs) in those overlapping with OL CHH-hyper DMRs in flanking sequences or gene body, relative to all genes. The hypergeometric test was used to infer statistical significance (*: P < 0.05; **: P < 1e^-10^). (*B*) CHH methylation distribution in genes of ovule-preferred and leaf-preferred genes in the leaf and ovule. Leaf-preferred and ovule-preferred genes were identified by both fold-change of gene expression (> 5-fold) and ANOVA test (< 0.01). (*C*) Relationships between CHH (left), CHG (middle) and CG (right) methylation and expression levels of genes in ovules, which were divided into four quartiles (high, red; mid-high, green; mid-low, blue; and low, yellow). (*D*) The relationship between siRNA density and the genes in the four quartiles as in (*C*). (*E*) Box plot showing siRNA abundance in promoter regions of ovule-preferred and leaf-preferred genes.

The higher methylation levels of most OL CHH-hyper DMRs in fibers than in ovules may suggest a role for increased methylation in fibers in gene expression. To test this, we extracted 18,107 OL CHH-hyper DMRs that showed higher methylation levels (cut-off value >0.05) in fibers than in ovules to examine effects of these DMRs on gene expression in fibers. In contrast to the up-regulation of genes in ovules, down-regulated genes in fibers relative to ovules were significantly enriched in the genes overlapping with these OL CHH-hyper DMRs in the upstream or downstream sequences, suggesting that further increasing methylation levels of some OL CHH-hyper DMRs in fibers represses nearby genes (Fig. 4A). For example, expression of GhMYB25L, a key factor for fiber development (Walford et al. 2011), was induced at the fiber initiation stage (0 DPA) but repressed in the elongation stage (Fig. 4B). Methylation levels in the flanking sequences of GhMyb25L increased in ovules at 0 DPA and continued to increase in fibers at 14 DPA. These data indicate that CHH hypermethylation in fibers regulates GhMYB25L expression during fiber development (Fig. 4B).

**Figure 4.**
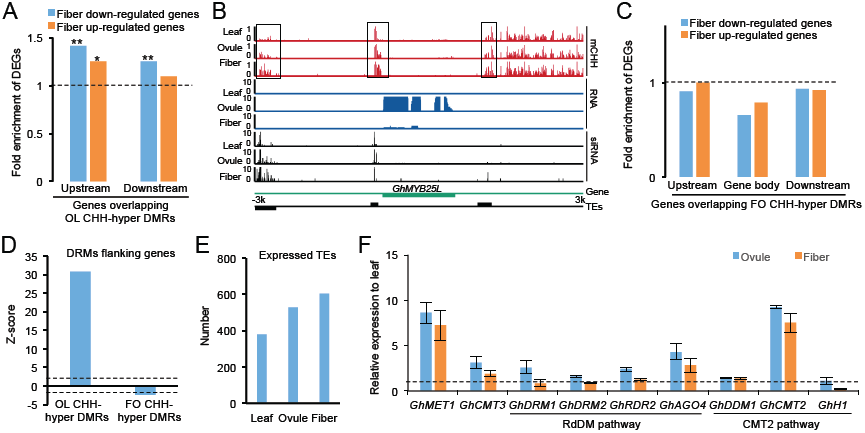
Influence of fiber-ovule (FO) CHH-hyper DMRs to gene expression and TE activity. (*A*) Fold-enrichment of DEGs between the fiber and ovule in the genes overlapping with OL CHH-hyper DMRs, which were further hypermethylated in fibers, relative to all genes. The hypergeometric test was used to infer statistical significance (*: P < 0.05; **: P < 1e^-10^). (*B*) An example of fiber CHH hypermethylation correlating with gene repression in the fiber. (*C*) fold-enrichment of DEGs between the fiber and ovule in the genes overlapping with FO CHH-hyper DMR in the gene body and flanking sequences, relative to all genes. (*D*) Enrichment (z-score) of DMRs in flanking sequences (1 kb) of genes. The dash line indicates the statistical significant level (P < 0.05). (*E*) Number of expressed TEs in the leaf, ovule and fiber. (*F*) Relative expression levels of genes related to DNA methylation in the leaf, ovule and fiber. Cotton *HISTONE H3* was used as internal control for qRT-PCR analysis. The dash line indicates the expression level that is normalized to the leaf.

### The functions of FO CHH hypermethylation

To our surprise, FO CHH-hyper DMRs were not significantly correlated with expression changes in fiber (Fig. 4C; Supplemental Table S4). This was partly because FO CHH-hyper DMRs tended to be excluded from the gene body (Fig. 2A) or from flanking sequences (1-kb) of the gene (Fig. 4D). Instead, FO CHH-hyper DMRs were enriched in TEs (Fig. 2A), and FO CHH hypermethylation was positively correlated with the TE density (Supplemental Fig. S1E). We predict that OL CHH hypermethylation mediates gene expression, while FO CHH hypermethylation represses TEs in heterochromatin. Indeed, RNA-seq data showed that more TEs were expressed in fibers than in ovules or leaves (Fig. 4E). These data indicate that CHH hypermethylation in fibers serve as a mechanism for repressing potential TE activation resulting from rapid fiber cell growth. As CHH methylation in heterochromatin is catalyzed by CMT2 through a DDM1-mediated process (Zemach et al. 2013; Stroud et al. 2014), we examined expression of *GhCMT2*, *GhDDM1* and other genes related to DNA methylation in leaves, ovules and fibers. Compared with leaves, *GhCMT2*, *GhCMT3* and *GhMET1* were up-regulated in ovules and fibers, and the genes involved in the RdDM pathway, including *DRM1*, *DMR2* and *RDR2* with an exception of *AGO4*, were up-regulated in ovules but not in fibers (Fig. 4D). However, cotton Histone H1, which is shown to impede the access of DNA methyltransferases to the heterochromatin (Wierzbicki and Jerzmanowski 2005; Zemach et al. 2013), was repressed only in the fiber but not in the ovule (Fig. 4F). The data suggest that up-regulation of *GhCMT2* and repression of cotton *HISTONE H1* may induce CHH hypermethylation in TEs and heterochromatin in fibers.

### Influence of DNA methylation to expression of homoeologous genes

Methylation in the gene body and promoter regions has different effects on expression of homoeologous genes. In the gene body, CG methlylation levels of most homoeologous genes were relatively equal (Fig. 5A) and did not correlate with homoeologous gene expression levels (Fig. 5B). However, we identified 533 pairs of A- and D-homoeologs that were differentially methylated at CHG and CHH sites, although non-CG methylation levels of most homoeologs were relatively low (Fig. 5A). The methylation levels of CHG or CHH methylation in the gene body were significantly anti-correlated with expression levels of A- and D-homoeologous alleles (Fig. 5B,C; Supplemental Table S5), indicating repression function of CHG and CHH methylation in expression variation of these homoeologous genes.

**Figure 5.**
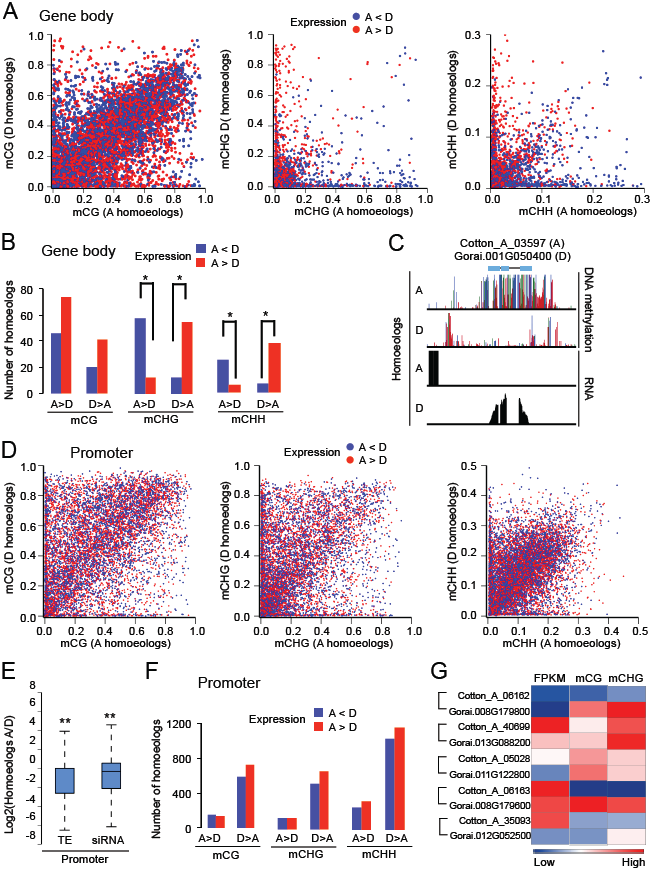
DNA methylation differences in homoeologous genes. (*A*) Pairwise plots of CG (left), CHG (middle), and CHH (right) methylation levels (Y-axis, D-homologs; X-axis, A-homoeologs) in the gene body for homoeologous genes. Blue and red dots indicate the expression level of an A-homoeolog that is lower and higher than that of the corresponding D-homoeolog, respectively. (*B*) Number of homoeologous genes that were differentially methylated in the gene body and showed expression differences (A < D, blue; A > D, red). The levels of CG (>0.4), CHG (>0.4), and CHH (>0.1) methylation difference and fold-changes of expression (>2-fold) between A- and D-homoeologs were used as cut-off values. (*C*) An example of hypermethylation in the gene body correlating with repression of the A-homoeolog. Colors indicate CG (green). CHG (blue), and CHH (red) methylation, and gene expression (black). (*D*) Pairwise plots of CG (left), CHG (middle), and CHH (right) methylation levels (Y-axis, D-homologs; X-axis, A-homoeologs) in promoter regions (2-kb) of homoeologous genes. Same colors were used as in (*A*). (*E*) Box plot showing more TEs and siRNAs in promoters of homoeologous genes derived from the D-subgenome than those from A-subgenome. (*F*) Number of homoeologous genes that were differentially methylated in CG (left), CHG (middle), and CHH (right) sites in promoter regions and showed expression differences (A<D, blue; A>D, red). Same cut-off values were used as in (*B*). (*G*) CG and CHG methylation levels in promoter regions of GhMYB25 family genes were anti-correlated with expression bias of A- and D-homoeologs. FPKM: Fragments per kilobase of transcripts per million mapped reads. Brackets indicate pairs of homoeologs.

In the promoter regions, although more TEs exist in the A-subgenome (68%) than in the D-subgenome (61%), more genes in the D-subgenome than in the A-subgenome had higher methylation levels in CG, CHG and CHH sites (Fig. 5D). This is consistent with more TEs and siRNAs that were distributed in promoter regions of D-homoeologs than those of A-homoeologs (P < 1e^-80^, Wilcoxon rank-sum test) (Fig. 5E). Homoeologous genes with higher CG or CHG methylation in promoter regions tended to be expressed at lower levels (Fig. 5F). For example, among 10 members of the *GhMYB25* family (homologs of *AtMYB25* or *AtMYB106*)(Machado et al. 2009; Walford et al. 2011), the D-homoeologs were methylated at higher levels in CG or CHG sites of promoter regions than the A-homoeologs and were also expressed at lower levels than the A-homoeologs (Fig. 5G). These data suggest a role for DNA methylation in the expression variation of homoeologous genes, which may contribute to fiber selection and improvement in the allotetraploid cotton.

## DISCUSSION

We found little variation of CG and CHG methylation among cotton tissues, which is consistent with the data in maize, showing more variable CG and CHG methylation levels among genotypes than among tissues (Eichten et al. 2013). Although CHH methylation is low and cannot be effectively detected by microarrays (Eichten et al. 2013), it links to biological functions. In *Arabidopsis* intraspecific hybrids, CHH methylation changes are associated with the parent-of-origin effect on circadian clock gene expression and biomass heterosis (Ng et al. 2014). CHH methylation is mainly produced by 24-nt siRNAs through the RdDM pathway (Law and Jacobsen 2010; Haag and Pikaard 2011). In cotton, 24-nt siRNAs were enriched in gene-rich regions, which is similar to that in maize (Gent et al. 2013) and sugar beet (Dohm et al. 2013) but different from that in *Arabidopsis* (Ha et al. 2009) and soybean (Song et al. 2013). Enrichment of siRNAs in euchromatic regions is inconsistent with the overall repeat abundance in the genome because the repeat amount is higher in soybean (˜59%) (Schmutz et al. 2010) than in sugar beet (˜42%) (Dohm et al. 2013), but much lower than in maize (˜85%) (Schnable et al. 2009). It is the TE property and location not the TE density that determines location of siRNA abundance and CHH methylation. These 24-nt siRNAs are derived from short TEs close to the genes, inducing CHH methylation flanking the genes, which is known as “CHH methylation island” in maize (Gent et al. 2013). However, in spite of similar siRNA distribution patterns between maize and cotton, CHH methylation patterns are different. In maize, siRNAs in gene-rich regions are associated with higher CHH methylation levels; but in cotton, CHH methylation levels were negatively correlated with the gene density and siRNA abundance (Fig. 1F). Furthermore, the percentage of methylcytosines in the CHH context is higher in cotton (˜7.7% in leaves) than in maize (˜5% in ears and shoots), which is in contrast to a higher TE density in maize than in cotton. These results indicate that in addition to CHH methylation that is produced by the RdDM pathway in cotton as in maize, cotton has another active pathway to generate CHH methylation.

This CHH hypermethylation of many TEs in fibers is likely mediated by CMT2 and DDM1 (Zemach et al. 2013). The genes overlapping with FO CHH-hyper DMRs were not enriched in the differentially expressed genes, which is consistent with the finding that non-CG methylation generated by CMT2 does not regulate protein-coding genes (Stroud et al. 2014). The CHH hypermethylation in cotton fibers is probably promoted by both induction of *GhCMT2* and repression of *HISTONE H1*. In ovules, *GhCMT2* is also induced but *HISTONE H1* is not up-regulated. As a result, CHH methylation was not enhanced in the heterochromatin of ovules, suggesting that chromatin changes by histone H1 is required for promoting CHH hypermethylation in heterochromatin between fibers and ovules. Fiber undergoes endoreduplication in early stages and rapid cell elongation and cellulose synthesis in later stages (Van’t Hof 1999; Wu et al. 2006). This dramatic change in the cellular environment (rapid elongation without cell division) of fibers could “elongate” or “loosen” chromatin structures in cell nucleus, leading to activation of TEs. Indeed, more TEs are active in fibers than in ovules and leaves (Fig. 2), and more siRNAs were generated in fibers than other tissues (Pang et al. 2009). The CHH hypermethylation induced in the fibers could function as a feedback mechanism to repress TEs. Consistent with the requirement of DNA methylation for cotton fiber development, inhibiting DNA methylation by aza-dC not only reduces fiber cell initials but also slows down fiber cell elongation (Fig. 1). A confirmatory experiment is to generate the transgenic plants that suppress CHH methylation and directly test the CHH methylation effect on fiber development.

Notably, the highly expressed genes are correlated with higher CHH methylation levels in promoter regions close (1-kb or less) to the transcription start site (TSS), which is unexpected but has been documented maize (Gent et al. 2013), and soybean (Song et al. 2013), and now in cotton (Fig. 3). One possibility is that TE activation enhances expression of nearby genes. Transcription initiates from TEs through Pol II or Pol IV and then spreads to nearby genes (Zheng et al. 2009; Haag and Pikaard 2011). While most TEs are transcriptionally silenced in plant genomes, some TEs could activate nearby genes, as reported in *Arabidopsis* (Wang and Warren 2010), rice (Naito et al. 2009), and wheat (Kashkush et al. 2003). If TE activation occurs prior to the transcription of nearby genes, TE expression should not be correlated with the distance between TEs and genes. However, more 24-nt siRNAs are present in the TEs closer to the genes (Fig. 1J), suggesting another possibility that gene expression induces activation of nearby TEs. This may contribute to positive correlation between CHH methylation in promoters and gene expression. The regions near the TSS are probably in open chromatin formation to allow active transcription of both genes by RNA polymerase II and short TEs by RNA polymerase IV, a homolog of Pol II (Herr et al. 2005; Onodera et al. 2005; Zheng et al. 2009). This leads to high abundance of small RNAs near the promoters and transcripts from corresponding genes, as observed in cotton, maize (Gent et al. 2013), and soybean (Song et al. 2013). The siRNAs can induce CHH methylation through the RdDM pathway. Consistent with this hypothesis, ovule up-regulated genes were enriched in those overlapping with OL CHH-hyper DMRs in flanking sequences. However, further CHH methylation of OL CHH-hyper DMRs in fibers represses nearby genes. This suggests that CHH methylation in promoters may act as a feedback mechanism to regulate these genes during ovule and fiber development. The spatiotemporal role of DNA methylation in expression changes of ovule- and fiber-related genes could also explain why overexpressing these genes may result in the unexpected outcome of fiber traits (Walford et al. 2011).

Finally, non-CG methylation in the gene body is associated with the expression bias of homoeologous genes in the allotetraploid cotton, providing the unique evidence for epigenetic regulation of non-additive expression of homoeologous genes in polyploid species (Liu and Wendel 2003; Chen 2007). Although *G. arboreum* (A-genome progenitor) (Li et al. 2014) contains more TEs than *G. raimondii* (D-genome progenitor) (Paterson et al. 2012), methylation levels and TE densities in promoters of many ovule- and fiber-related genes are higher in the D-homoeologs than in the A-homoeologs. Consequently, A-homoeologs of ovule- and fiber-related genes are expressed at higher levels than their D-homoeologs. This could reflect an evolutionary difference between fiber-bearing A-genome progenitors and nearly fiberless D-genome progenitors. Together, these results will advance our understanding of the biological significance of CHH methylation in developmental regulation and intergenomic interactions, which could be translated into genomic and biotechnological improvement of in polyploid plants, including most important crops that provide us with food (wheat), fiber (cotton), and oil (canola).

## METHODS

### Plant material

*Gossypium hirsutum* acc. TM-1 grew in the greenhouse at The University of Texas at Austin. Leaves and bolls (0 and 14 DPA) were harvested with three biological replications. Ovules and fibers were carefully dissected from cotton bolls at 14 DPA and immediately frozen in liquid nitrogen for RNA and DNA preparation.

### Small RNA-seq library construction

Total RNA was isolated from cotton leaves, ovules at 0 DPA and fibers at 14 DPA using Plant RNA Reagent (Life Technologies). By polyacrylamide gel electrophoresis, RNAs corresponding to ˜15 to 30-nt in length from 20 μg total RNA were excised and eluted from the gel using 0.3 M NaCl. Small RNAs were precipitated using ethanol and dissolved in 6 μl RNase free water. Small RNA-seq libraries with three biological replicates were constructed using NEBNext^®^ Multiplex Small RNA Library Prep Set (NEB, Ipswich, Massachusetts) according to the manufacturer’s instructions. Small RNA-seq libraries were single-end sequenced for 50 cycles.

### mRNA-seq library construction

After DNase treatment, total RNA (˜5 μg) was subjected to construct strand-specific mRNA-seq libraries with two biological replications using NEBNext^®^ Ultra^™^ Directional RNA Library Prep Kit (NEB, Ipswich, Massachusetts) according to the manufacturer’s instructions. mRNA-seq libraries were single-end sequenced for 150 cycles.

### MethylC-seq library construction

Genomic DNA was isolated from cotton leaves, ovules at 0 DPA and 14 DPA, and fibers at 14 DPA using CTAB method (Mei et al. 2004). Total genomic DNA (˜5 μg) was fragmented to 100-1000 bp using Bioruptor (Diagenode, Denville, New Jersey). End repair (NEBNext^®^ End Repair Module) was performed on the DNA fragment by adding an ‘A’ base to the 3’end (NEBNext^®^ dA-Tailing Module), and the resulting DNA fragment was ligated to the methylated DNA adapter (NEXTflex^™^ DNA Barcodes, Bioo Scientific, Austin, Texas). The adapter-ligated DNA of 200–400 bp was purified using AMPure beads (Beckman Coulter, Brea, California), followed by sodium bisulfite conversion using MethylCode^™^ Bisulfite Conversion Kit (Life Technologies, Foster City, California). The bisulfite-converted DNA was amplified by 12 cycles of PCR using LongAmp^®^ *Taq* DNA Polymerase (NEB, Ipswich, Massachusetts) and purified using AMPure beads (Beckman Coulter, Brea, California). The Paired-End sequencing of the MethylC-seq libraries was performed for 101 cycles.

### qRT-PCR

After DNase treatment, total RNA (2 μg) was used to produce first-strand cDNA with the Omniscript RT Kit (Qiagen, Valencia, California). The cDNA was used as the template for qRT– PCR using FastStart Universal SYBR Green Master (Roche, Indianapolis, India). The reaction was run on the LightCycler^®^ 96 System (Roche, Pleasanton, California). The relative expression level was quantified using internal control cotton *HISTONE H3*.

### Reference genome generation

Genome sequences of *G. arboreum* (Li et al. 2014) and *G. raimondii* were respectively downloaded from http://cgp.genomics.org.cn and http://phytozome.jgi.doe.gov (Paterson et al. 2012), respectively. DNA sequencing data of *G. hirsutum* cv. Acala Maxxa (SRR617482) was downloaded from NCBI (Paterson et al. 2012). The reads were mapped to genome sequences of *G. arboreum* and *G. raimondii* using Bowtie2 with default parameter and only mismatches with high base quality score (>=30) were used for variant calling (Langmead and Salzberg 2012). Calling variants was performed by SAMtools (Li et al. 2009). A SNP was identified when the identity was more than 90% and read depth was over 10. We substituted each SNP in genome sequence of *G. arboreum* and *G. raimondii* to construct *G. hirsutum* genome, which was used as reference genome for aligning all high-throughput sequencing data in this study.

### Small RNA-seq data analysis

After adapter clipping, small RNA-seq reads (18-30 nt) were mapped to reference genome using Bowtie2 allowing no mismatches. Multi-mapped small RNA-seq reads were evenly weighted and assigned to all locations as previously described method (Lu et al. 2012).

#### mRNA-seq data analysis

Gene annotation and TE classification of *G. arboreum* and *G. raimondii* were downloaded from http://cgp.genomics.org.cn and http://phytozome.jgi.doe.gov, respectively. mRNA-seq reads were mapped to reference genome using Tophat (Trapnell et al. 2009). Uniquely mapped reads were extracted and analyzed by Cufflinks to determine gene and TE abundance using annotated genes and TEs (Roberts et al. 2011). The up-regulated or down-regulated genes were identified through double-filtering methods using the fold-change (>2-fold) and ANOVA tests (P < 0.01).

#### Identification of methylated cytosines

We applied Bismarck software to align reads to reference genome with default parameters (Krueger and Andrews 2011). In brief, the first 75 bases of unmapped reads were extracted and realigned to the genome. Only reads mapped to the unique sites were retained for further analysis. Reads mapped to the same site were collapsed into a single consensus read to reduce clonal bias. For each cytosine, the binomial distribution was used to identify whether this cytosine was methylated. The probability *p* in binomial distribution B (n, *p*) was referred to bisulfite conversion failure rate. The number of trials (n) in the binomial distribution was the read depth. Only the cytosines covered by at least three reads in all compared tissues were considered for further analysis. Cytosines with P-values below 1e^−5^ were identified as methylcytosines.

#### Differentially methylated regions

Differentially methylated regions (DMRs) for CHH methylation were identified using 100-bp sliding-windows. Mean methylation level was calculated for each window (Schultz et al. 2012). Windows containing at least eight cytosines in the CHH context covered by at least three reads were selected for identifying DMRs. ANOVA test using 2 biological replications (*P* < 0.05) and difference of methylation levels (cut-off value > 0.1 between two compared samples) were used to determine CHH DMRs between two compared tissues. The cut-off value was set at 0.05 for the comparison between ovule and fiber methylation levels in OL CHH-hyper DMRs.

#### Homoeologous genes

Protein sequences of *G. arboreum* were aligned to protein sequences of G. raimondii using blastp with an E-value less than 1e^-10^. Alignment information was used by MCScanx to identify homoeologous genes (Wang et al. 2012) (Score > 2000 and E-value < 1e^-10^).

#### Z-score for enrichment of DMRs flanking genes

We first randomly extracted 1000 of 1-kb regions in the genome for 500 times. Percentage of DMR-overlapping regions in 1000 random regions was calculated at each time. Z-scores were calculated as (*x-μ*)/*σ* where x was percentage of DMR-flanking genes in all genes, *μ* and *σ* were mean and standard deviation of percentages of DMR-overlapping regions in random loci.

#### Cotton ovule in vitro culture and 5-aza-dC treatment

Ovules were removed from flower buds at -3 or 0 DPA. The ovules were sterilized with 75% alcohol, washed with autoclaved water, and cultured in Beasley-Ting (BT) liquid media (20 ml) containing 5 μM indole-3-acetic acid (IAA) and 0.5 μM gibberellic acid3 (GA3) at 30°C for 4-14 days in dark (Beasley and Ting 1974; Kim and Triplett 2001). Treatment of 5-aza-2’-deoxycytidine (aza-dC, 10 mg/L) was applied in the BT liquid media.

#### Data access

All raw data have been submitted to NCBI GEO database under accession number GSE61774.

## Acknowledgements

We thank the Genomic Sequencing and Analysis Facility at The University of Texas at Austin for sequencing methylomes, RNA, and small RNA libraries using HiSeq 2500. This work is supported by the grants from the National Science Foundation Plant Genome Research Program (IOS1025947) and the Cotton Incorporated (07-161).

### Author contributions

QS and ZJC conceived and designed the project; QS performed the experiments; XG provided materials and reagents; QS and ZJC analyzed data and wrote the paper.

